# Towards high throughput in-field detection and quantification of wheat foliar diseases with deep learning

**DOI:** 10.1101/2024.05.10.593608

**Authors:** Radek Zenkl, Bruce A. McDonald, Achim Walter, Jonas Anderegg

## Abstract

Reliable, quantitative information on the presence and severity of crop diseases is critical for site-specific crop management and resistance breeding. Successful analysis of leaves under naturally variable lighting, presenting multiple disorders, and across phenological stages is a critical step towards high-throughput disease assessments directly in the field.

Here, we present a dataset comprising 422 high resolution images of flattened leaves captured under variable outdoor lighting with polygon annotations of leaves, leaf necrosis and insect damage as well as point annotations of Septoria tritici blotch (STB) fruiting bodies (pycnidia) and rust pustules. Based on this dataset, we demonstrate the capability of deep learning for keypoint detection of pycnidia (*F* 1 = 0.76) and rust pustules (*F* 1 = 0.77) combined with semantic segmentation of leaves (*IoU* = 0.96), leaf necrosis (*IoU* = 0.77) and insect damage(*IoU* = 0.69) to reliably detect and quantify the presence of STB, leaf rusts, and insect damage under natural outdoor conditions. An analysis of intra- and inter-annotator agreement on selected images demonstrated that the proposed method achieved a performance close to that of annotators in the majority of the scenarios.

We validated the generalization capabilities of the proposed method by testing it on images of unstructured canopies acquired directly in the field and without manual interaction with single leaves. The corresponding imaging procedure can be adapted to support automated data acquisition. Model predictions were in good agreement with visual assessments of in-focus regions in these images, despite the presence of new challenges such as variable orientation of leaves and more complex lighting. This underscores the principle feasibility of diagnosing and quantifying the severity of foliar diseases under field conditions using the proposed imaging setup and image processing methods. By demonstrating the ability to diagnose and quantify the severity of multiple diseases in highly natural complex scenarios, we lay out the groundwork for a significantly more efficient, non-invasive in-field analysis of foliar diseases that can support resistance breeding and the implementation of core principles of precision agriculture.

## 2 Introduction

Modern mitigation of plant diseases in agriculture relies heavily on pesticides and resistant plant varieties [22]. Unfortunately, pathogens can adapt to both host resistance and pesticides used to control them [21] which poses a recurrent threat to global food security. Resistant plant varieties typically contain a combination of resistance genes or even just a single resistance gene [25] [13] which can prompt boom-and-bust cycles where the pathogen overcomes the resistance mechanism. Cases of pathogens overcoming resistance genes within only a few years of their introduction have been well documented [16] [41] [47].

One of the tools that modern breeders commonly employ is qualitative, major resistance genes [36] [38]. The presence of a major resistance gene can be readily ascertained using inoculation assays under controlled conditions. Once a major resistance gene has been identified, isolated and introgressed into commercial germplasm, it can be deployed in the field with a high confidence of success because the underlying mechanism involves the direct interaction between the pathogen and the host. In the case of Quantitative Resistance (QR), multiple genes with minor effects typically contribute to resistance [36] [38]. The mechanisms underlying QR can have varying degrees of complexity. In contrast to gene-for-gene mechanisms, hosts typically do not exhibit the same boom-and-bust cycles but rather a gradual decrease in QR efficiency as pathogens slowly adapt to QR. Thereby leading to a much longer lasting resistance effect [13] [36] [38] [37].

The search for QR requires a precise and quantitative assessment of the disease intensity when the host is interacting with pathogen populations in a natural environment. These interactions vary depending on the crop growth stage, many environmental factors, some of which some can be influenced by field management practices, existing disease pressure, and co-infections with other diseases, amongst other factors [35] [44] [1]. It is therefore not feasible to evaluate QR reliably under controlled conditions [45]. Moreover assessments of disease escape and disease tolerance are determined based largely on yield or require realistic canopy structure and plant morphology for reliable assessment, also calling for field-based evaluation of disease.

Since QR leads to a reduction rather than an absence of the disease symptoms, evaluating symptoms at a single time point does not describe the full impact of QR because it has the potential to alter epidemic progression through many effects, including by reducing pathogen reproduction. Therefore, it is important to measure the effect of QR at the right time [23]. In addition, the tolerance of many host plants to diseases is influenced by the developmental stage at the time point of exposure to the pathogen [49], so when assessing the potential yield loss it is necessary to describe the extent of the epidemics with respect to the phenological stage of the crop [10]. Repeated measurements during a growing season not only ensure that the cultivars are assessed at the appropriate time, but also allow modelling of the epidemics progression over time and reduce measurement variance through having multiple observations.

A lack of accurate phenotypic data is the main reason for the current under-utilization of QR in breeding programs [26] [38] [50] [15]. The gain of a breeding program with respect to QR scales directly with the experiment size as larger experiments allow for a larger volume of genetic material to be tested [9] [26]. Unfortunately, evaluating the necessary field experiments is associated with an enormous amount of phenotypic variance coming from many sources including weather patterns, spatial dependence, and co-exposure to other stresses, which cannot be controlled by breeders. According to [42] in a multi environment field trial for disease resistance in wheat varieties, the variance coming from *environment × genotype* accounted for more than 50% of the overall genotypic variance. This underlines the need for new assessment methodologies capable of evaluating large volumes of data to separate the effects of QR from other factors while processing large amounts of genetic material.

The goal of this work was to investigate Septoria Tritici Blotch (STB), the most damaging wheat disease in Europe, caused by the fungal pathogen *Zymoseptoria tritici*, as a model disease for acquiring quantitative phenotype dataset under field conditions. More specifically, we aim to develop a high throughput method capable of diagnosis and accurate assessment of STB in uncontrolled outdoor lighting during co-infection with other naturally occurring diseases. The underlying task consists of assessing the Percentage of Leaf Area Covered by Lesions (PLACL) as a quantification of the host damage and the pycnidia density as a measure of reproductive potential for STB. The latter is especially important as STB epidemics are driven by secondary inoculum involving asexual spores originating from pycnidia [53] which determine the overall damage and resulting yield loss potential. The measures of PLACL and pycnidia are largely independent in natural field infections and thus both need to be measured [30]. Several different approaches in terms of sensor modality, resolution and throughput have already been explored. The oldest and still most widely used assessment relies on visual scoring by trained personnel, where the PLACL and pycnidia density are each assigned to a class based on a visually estimated severity [40] [43] [20] [14]. As these measurements are time intensive, significant efforts have been invested into achieving higher throughput. The developed methods can be grouped based on the level of detail that they achieve, with many methods sacrificing the ability to resolve individual pycnidia to achieve higher throughput by using RGB or multispectral imagery with lower physical resolution [48] [8] [11] or one dimensional spectral measurements on a small plot basis [52] [3] [6]. On the other side of the detail spectrum, methods utilizing very high resolution of more than 0.01 mm per pixel, were developed to detect individual pycnidia in addition to the necrotic lesions [30] [45] [34]. However, these latter methods require invasive, tedious sample preparation to capture high resolution images, greatly limiting the potential throughput. None of the mentioned methods is capable of handling naturally occurring co-infections due to missing data or an inability to distinguish among the diseases due to the underlying evaluation method. Novel methods increasingly utilize data driven deep learning methods [34] [8] instead of color thresholding and visual indices [30] [45] [48]. To the best of our knowledge, no prior work has attempted large scale STB assessments that can resolve individual pycnidia under natural field conditions. We believe that conducting STB phenotyping under natural field conditions represents the next key step to improving qunatitatice STB resistance in breeding programs. Our aim is to achieve throughput without sacrificing quality in order to enable scoring of STB resistance in non-invasive manner or even in a fully autonomous manner without any human input.

## 3 Materials and Methods

### 3.1 Data Acquisition

To achieve a representative sample of STB symptoms under natural conditions, we conducted a field experiment using a set of 16 wheat genotypes exposed to various treatments, including inoculation with a mixture of *Zymoseptoria tritici* strains. Wheat cultivars were chosen to maximise variation in terms of morphology and STB susceptibility based on data from [2] and [30]. The key selection factors were planophile and erectophile leaves, high and low levels of flag leaf glaucousness and degree of STB resistance or susceptibility. A mixture of 10 Swiss *Z. Tritici* isolates named 1A5, 1E4, 3A1, 3A8, 3B2, 3B8, 3D1, 3D7, 3G3, 3F5 were selected to maximise symptom diversity based on data from [19]. Three treatments consisting of different combinations of fungicide applications and artificial inoculations were applied. For more extensive information about the experimental design see [8].

One of the key challenges in acquiring highly detailed images in an outdoor environment is the trade-off between maximizing the scanned area whilst guaranteeing sufficient ground sampling distance. The limiting factor in our task is the appropriate imaging of pycnidia which are around 0.1 mm in diameter. For very small features such as pycnidia, a high physical resolution of the imaging setup is required to achieve sufficient resolution. Our data was collected using a full-frame mirrorless digital camera (EOS R5, Canon Inc., Tokyo, Japan; 45 megapixel, 36*×*24 mm sensor) combined with a macro lens (RF 35 mm f/1.8 IS Macro STM, Canon Inc., Tokyo, Japan). The camera was mounted on a custom-made stand to ensure a consistent working distance of 23cm. Such close up imaging with a high-resolution sensor led to challenging behaviour in terms of exposure and blur, due to the small size of a pixel on the sensor, in our case 4.39µm. Our imaging setup resulted in a physical resolution of approximately 0.03 mm/px. For comparison, the earlier flatbed scanning technique [45] using 1200dpi provided a resolution of approximately 0.02 mm/px.

During the image acquisition, the F-stop was fixed at 10 to provide a reasonable trade off between depth-of-field, amount of incoming light and lens diffraction. Concerning the sensor gain (ISO), the lowest possible values are preferred because the sensor gain amplifies the sensor noise as well. In the scope of resolving small objects, the resulting size of noise peaks operates on a similar scale as pycnidia due to the effect of de-bayering where one noisy pixel influences its neighbours as well. ISO was kept under 3200 to prevent high levels of background noise. The upper bound for ISO was determined experimentally by observing the noise dynamics with respect to resulting contrast between pycnidia and background noise. The F-stop and ISO settings led to a typical exposure times between 1/100 and 1/1000 sec depending on the lighting conditions during acquisition.

Two flag leaves per plot were detached and photographed outdoors under both diffuse and direct lighting when possible. The leaves were fixed onto a blue background plate by gently pressing them onto a temporary glue. This process was repeated for three separate time points leading to 432 leaves imaged mostly under both diffuse and direct lighting. As the whole imaging process was conducted in the field, direct lighting was available only under clear skies. Diffuse lighting, on the other hand, was achieved by the operator casting a shadow over the imaging setup. The scanned leaves were not moved between the diffuse and direct light images, so the images were identical except for the different forms of lighting. After the field imaging, the leaves were flattened and additional flatbed scanner images were obtained according to the protocol in Stewart et al. [45].

**Figure 1:**
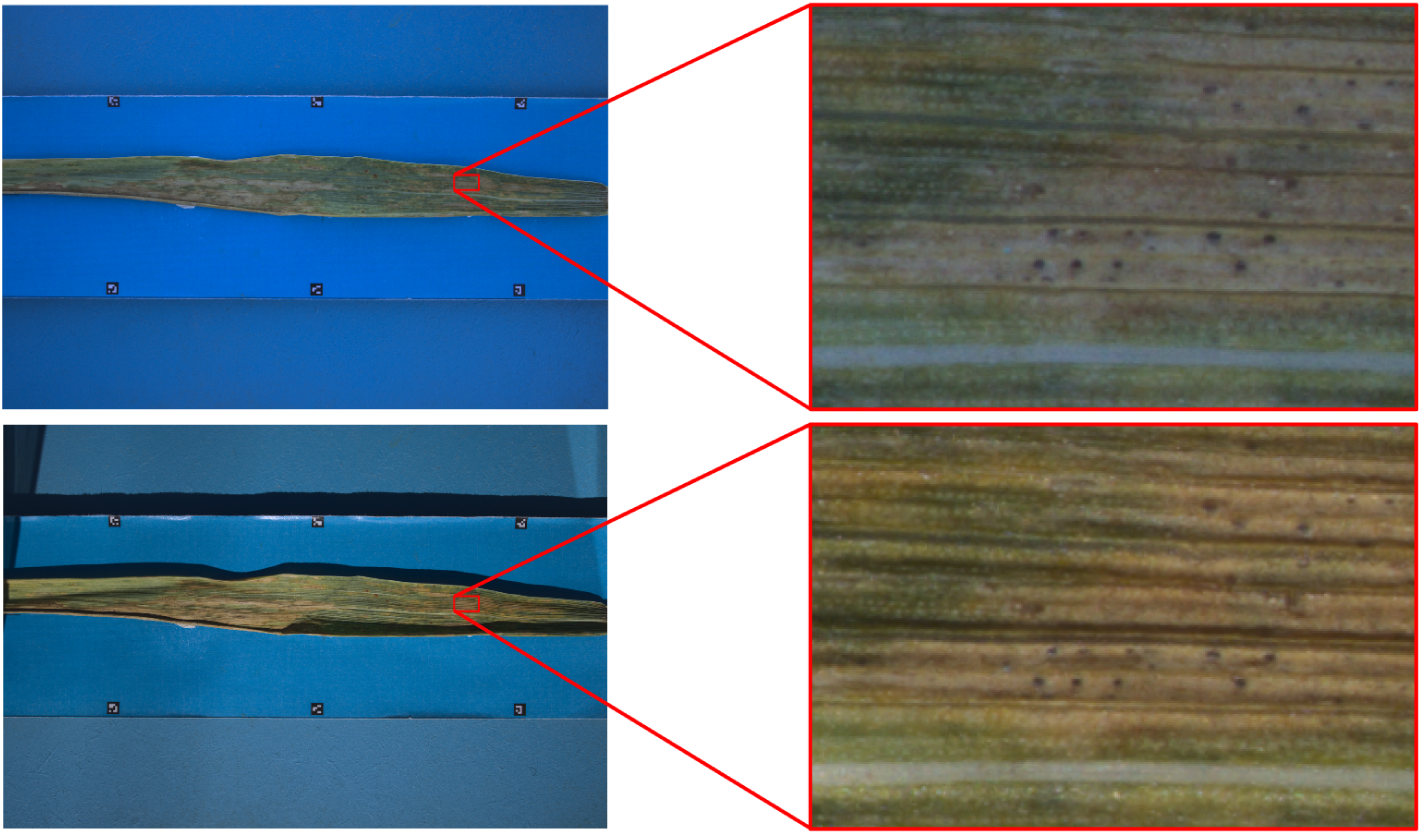
Sample image pair from the described imaging setup. The top image was taken under diffuse lighting. The bottom image was taken under direct sunlight.

### 3.2 From Images to Deep Learning Dataset

Each collected image was cropped to form 8 smaller patches of 1024*×*1024 px. From the resulting set of cropped images a subset of 422 images was selected at random for annotation, forming the Eschikon Foliar Disease (EFD) dataset.

Completely healthy or completely necrotic samples were removed because leaves were not useful for symptoms training, whilst completely necrotic leaves tended to be senescent rather than heavily diseased. All images were annotated using the CVAT [17]. The annotated classes were: necrotic tissue, leaf, insect damage, STB pycnidia and leaf rust pustules. To simplify the annotation process, STB pycnidia and leaf rust pustules were annotated as points placed at the object’s centroid. The remaining classes were annotated on a pixel level using polygons. Annotated images covered a wide range of scenarios (see Figure 2) due to different lighting conditions, varying symptoms, different symptom severities, the occurrence of chlorosis and different host genotypes at different phenological stages as described in subsection 3.1.

The semantic meaning of the annotations leaves room for potential mistakes due to overlapping regions such as a pycnidium on top of a necrotic lesion which itself is located on top of a leaf. To ensure reproducible results, the annotated polygons were exported as pixel-level segmentation masks. The individual masks from the polygon annotations were stacked using the following order where the previous labels are overwritten by the ones with higher priority. First, the polygon annotations are converted: the base layer is the leaf, followed by necrosis and insect damage. Regarding the point annotations for pycnidia and rust pustules, the point annotations are denoted in a YOLO-Pose dataset format^1^. The annotations of keypoints were extended by estimated bounding boxes of size 8*×*8px for pycnidia and 32*×*32px for rust pustules. This combination of formats should ensure repeatable and flexible use of the dataset.

#### 3.2.1 Label Uncertainties

Utilizing manual human annotations is the gold standard for creating the ground truth data for training and validation. Unfortunately, outdoor-grown leaf samples exhibit many ambiguities due to ill-defined symptom boundaries, inconclusive diagnosis of symptoms, and occasional images with lower quality. To put performance metrics into perspective, we quantified the uncertainty of the labels by conducting multiple independent annotations on the same images and investigating the resulting differences. For this, we selected 10 images from the validation set where we annotated lesions and pycnidia. Each of five annotators annotated each image twice, leading to a set of 10 labelled sets per image. This data enabled a quantification of the inter-annotator and intra-annotator reliability. [12] By utilizing a one-vs-rest crossvalidation, we computed IoU for lesions and F1 scores for pycnidia, where detection of pycnidia is considered correct when its placement is within 5px of the reference. Finally, to put the annotators’ performance into perspective, we compared the performance of the proposed method with respect to the annotator statistics by computing the metrics for each annotation attempt separately.

**Figure 2:**
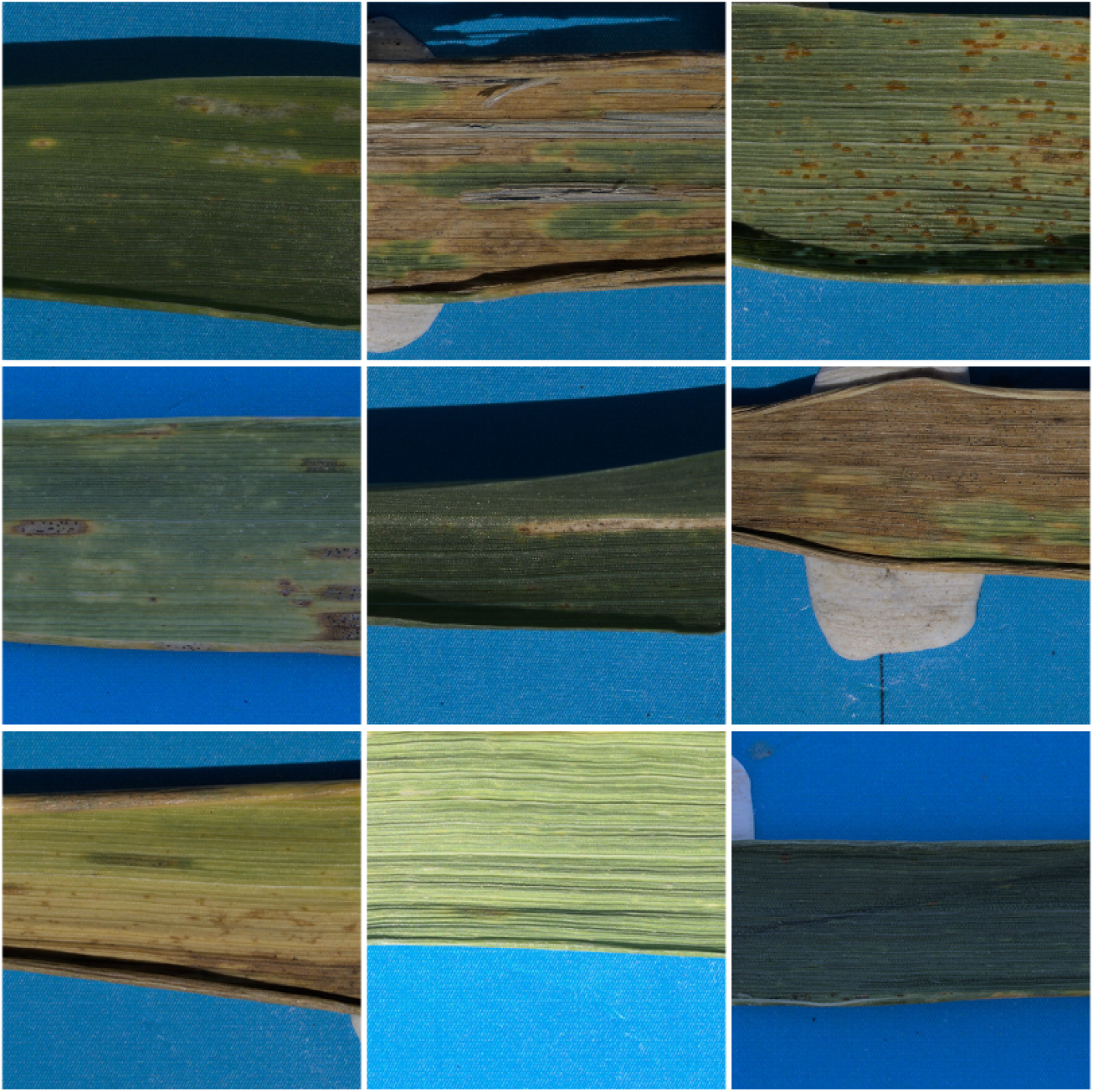
Samples from the proposed dataset which demonstrate the diversity of the images.

### 3.3 Image Processing Method

The overall goal of the image processing method can be summarized as keypoint detection and semantic segmentation. For the sake of simplicity both tasks are handled separately by individual models. For the task of semantic segmentation, a SegFormer backbone network [51] was used in combination with the FPN decoder [31]. The utilization of self-attention in this novel architecture allows for a significantly wider effective receptive field compared to convolutional neural networks [51]. In the context of disease diagnosis, this feature can be advantageous when context from a distant region is needed for diagnosis. Especially under changing light it might be beneficial to be able to get a reference for chlorosis in order to assess whether a given segment is already necrotic or not. The proposed combination of backbone and head enables inference on arbitrary resolution which can differ from the training set. In our case we can train on diverse small image crops while the inference can be done on full resolution images without a loss of performance. In addition, the SegFormer Backbone is significantly more robust to distortions such as blur, or noise which naturally occur during outdoor computer vision tasks [51]. Keypoint Detection is achieved by using the YOLOv8 Pose model [29] which utilizes an anchor-free detection head. This allows for effortless change of input size resolution without the need for tuning of anchor-boxes. During training the extensive data augmentation pipeline from ultralytics has been used.

Regarding the semantic segmentation, the very basis of hyperparameter tuning was the selection of an appropriate backbone network, responsible for the feature extraction. The palette of tested models was limited to one model family. The rationale behind this is to limit the extensiveness of the resulting hyperparameter search space in favor of exploring various depths of the models from the same model family rather then testing completely different model families. For semantic segmentation the tested models are of the SegFormer Backbone [51] architecture at different depths. The selection of a backbone’s depth has a major impact on the overall performance of the model in terms of performance, runtime and GPU Memory consumption. All tested models were trained with the Adam optimizer, batch size 4, images with full resolution of 1024*×*1024px for 50 epochs. All semantic segmentation models were trained using the cross entropy loss in a finetuning manner starting from Imagenet [18] pretrained weights while using the FPN [31] head. The results for semantic segmentation are reported in Table 1. The same approach was also applied to the keypoint detection network which uses the YOLOv8-pose architecture which offers different depths of the backbone [29]. The keypoint detection models were pretrained on COCO 2017 keypoint dataset [33]. For the training, we used YOLOv8’s default data augmentation.

Reported numbers correspond to the randomly sampled validation split of the dataset leading to 354 training images (80%) and 88 validation images (20%). The same split was applied to both, semantic segmentation and keypoint detection.

#### 3.3.1 Generalization Capabilities

Due to the small dataset size, additional datasets from other domains were evaluated to quantify the robustness and transferability of the trained models. Sections 4.3.1 and 4.3.2 illustrate the performance based on very different scenarios. The inference on images captured with flatbed scanners showcase the capabilities of dealing with another imaging sensor (LiDE 400, Canon Inc., Tokyo, Japan), a different background and artificial lighting. This imaging setup corresponds to the method proposed by [45]. Following the analysis protocol a white background color is used on the images. More importantly, the scanned leaves were detached from wheat plants and pressed flat prior to the imaging, hence the visual appearance of the leaves has been altered compared to the EFD images. As a second test domain, field images of the canopy were selected. This corresponds to a scenario much closer to the target application. However it contains several different aspects, not covered by the proposed dataset. The canopy images are acquired from a side sngle and cover approximately the two topmost leaf layers. Besides the expected leaves, the images contain ears, stems and no clear background. The images suffer from large amounts of blur for objects that are not in the focus plane. In contrast to the previous two setups, the leaves are freely oriented in space rather than fixed onto a flat surface. Finally, the lighting is much more complex due to occluding plant organs and leaves being semi-transparent, especially when illuminated from the back.

## 4 Results

### 4.1 Model Performance

To achieve for an optimal encoder depth, we benchmarked various depths of SegFormer [51] for semantic segmentation (Table 1) and of YOLOv8 [29] for keypoint detection (Table 2). The evaluation in Table 1 indicated that an increase in backbone complexity led to an improvement in performance. The more complex backbones mitb1 and mitb2 outperformed mitb0 mostly due to the large gap in IoU of insect damage. Comparing the performance of mitb1 and mitb2 showed only a minor difference in IoU of lesions but a modest difference in performance regarding insect damage in favor of the less complex mitb1. This left mitb1 as the best candidate for a backbone offering best performance whilst avoiding unnecessary computational complexity. Generally, a large discrepancy in performance of individual classes was observed, where the class “leaf” always achieved a very high score whilst “lesions” and “insect damage” produced much lower scores. This effect can be attributed to a large imbalance in the number of pixels assigned to the different labels, where the classes lesion and insect damage are under-represented compared to leaves. Under-represented classes provide less information for training which can lead to worse performance. In addition, smaller objects have higher boundary to area ratios making the achievement of high metric scores more challenging.

Similar behaviour in performance and model complexity can be observed in keypoint detection. Optimal performance was achieved with model YOLOv8m-pose, which performs on par with its more complex counterpart YOLOv8l-pose, whilst outperforming the less complex variants (Table 2). Interestingly, pycnidia exhibit a similar F1 Score as rust pustules (*F* 1*_pycnidia_* = 0.76 and *F* 1*_rust_* = 0.77), whilst there were approximately three times more instances of pycndia than rust pustules. Both pycnidia and rust pustules are confused with background but not with one another, leading to high confidence of disease assignment to the symptoms (see Supplementary Table A1).

**Table 1:** Overview of the performance of different backbones for Semantic Segmentation.

| Model Backbone | # param. | mIoU | IoU Leaf | IoU Lesion | IoU Insect Damage |
| --- | --- | --- | --- | --- | --- |
| mitb0 | 3.7M | 0.76 | 0.96 | 0.77 | 0.53 |
| mitb1 | 14.0M | 0.81 | 0.96 | 0.77 | 0.69 |
| mitb2 | 25.4M | 0.79 | 0.95 | 0.76 | 0.64 |

**Table 2:** Overview of the performance of different YOLOv8 models for keypoint detection.

| Model | # param. | mAP50 | mAP50-95 | F1 score |
| --- | --- | --- | --- | --- |
| YOLOv8n-pose | 3.3M | 0.78 | 0.77 | 0.72 |
| YOLOv8s-pose | 11.6M | 0.80 | 0.79 | 0.75 |
| YOLOv8m-pose | 26.4M | 0.82 | 0.81 | 0.76 |
| YOLOv8l-pose | 44.4M | 0.82 | 0.81 | 0.76 |

### 4.2 Data Quality and Labels Uncertainty

The annotator agreement and the resulting metrics seem to be consistent for the majority of the images with both the lesion IoU mean and pycnidia F1 score mean achieving values around 0.85 with typically lower values for lesion Iou compared to pycnidia F1 scores (see Figure 3). To quantify the potential for improvement, differences between annotation score median and prediction scores median was computed. For Lesion IoU the mean error was 0.05 and the error standard deviation was 0.03. Similarly, for Pycnidia F1 score the mean error was 0.03 and standard deviation was 0.04.

Generally, two types of ambiguities were discovered: 1) perturbations along the annotation contours; 2) entire regions which were assigned a different class. The former arose due to annotation inaccuracies or ill-defined lesion edges (see Figure 3.b.2 and Figure 3.b.3). After considering conflicting annotations of entire regions, a negative impact due to early or late stage rust symptoms could be identified (see e.g. Figure 3.b.5). In addition, damaged regions without pycnidia, especially on the rolled up bottom part of the leaf, showed lower annotator agreement (see e.g. Figure 3.a.3). A similar behavior can be observed for newly forming lesions which partially contain pycnidia whilst not being clearly necrotic (see Figure 3.a.4).

#### 4.2.1 Diseases Evaluation

The most intuitive parameter of the assessment of the disease analysis is its performance with respect to the indicators that breeding programs can utilize. In the case of STB analysis, this corresponds to the Percent Leaf Area Covered by Lesions (PLACL) and the number or density of pycnidia on leaves. Our proposed method achieved Mean Squared Error (MSE) of 4.0e-3 for PLACL and F1 Score of 0.76 for pycnidia. Figure 4 shows a success case under diffuse lighting and a failure case under direct lighting. Visual inspection of Figure 4 a) reveals that the vast majority of pycnidia and rust pustules were correctly predicted. The necrotic lesion is correctly separated from healthy leaf tissue and chlorotic regions. In Figure 4 b) the method achieves a solid performance in the upper part of the leaf, whilst the negative impact of the hard shadows can be observed based on the inconsistent boundaries of lesions in the bottom part of the leaf.

**Figure 3:**
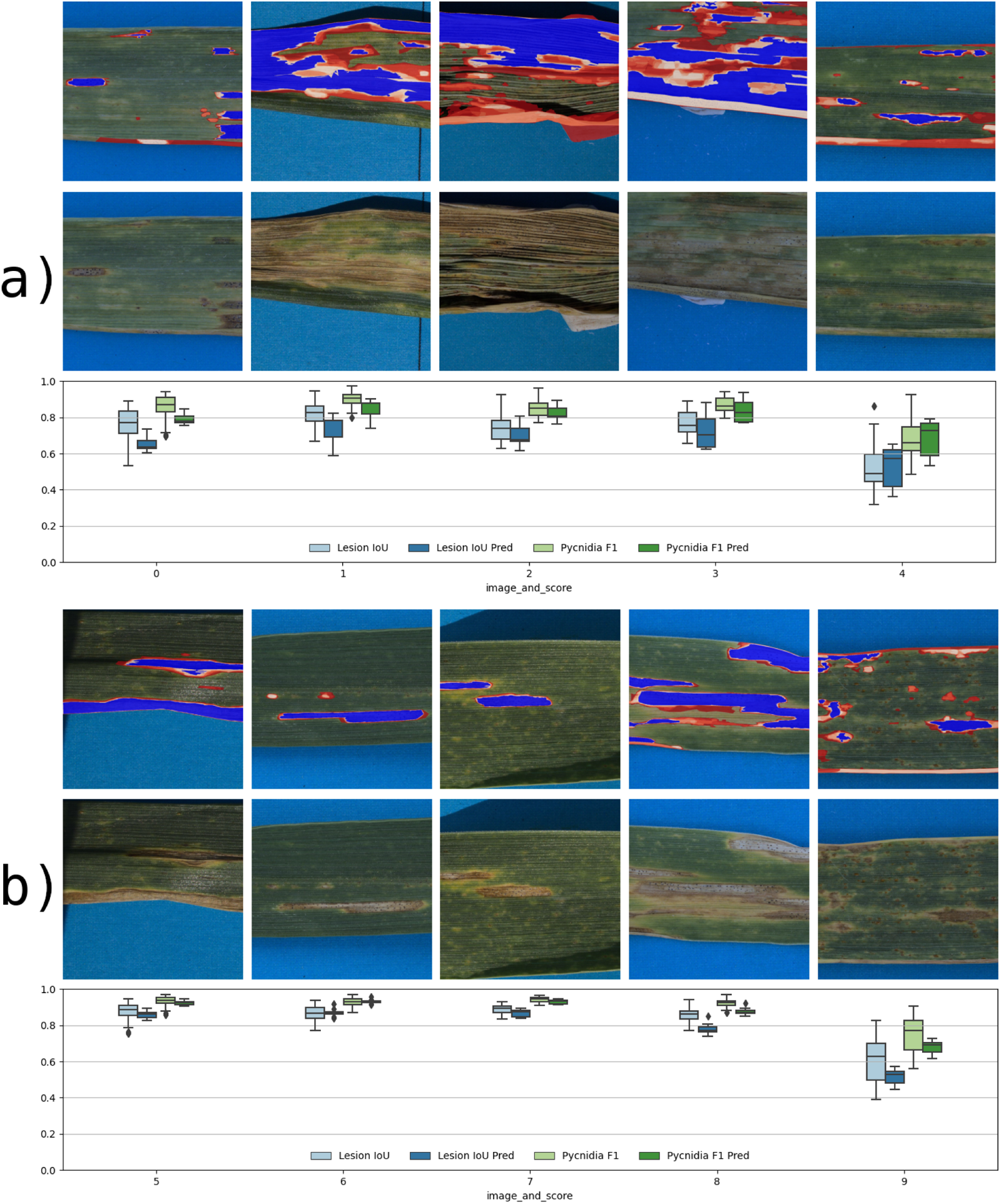
Selected 10 images organized in panels a) and b). The top row of each panel corresponds to an annotation overlay where blue color denotes a 100% agreement of all annotation runs. The disagreement among the annotation attempts is denoted in shades red where white denotes high agreement (many annotator attempts classified the region as lesion) and saturated red denotes low agreement (e.g. single annotator attempt classified the region as lesion). The middle row of the panels shows the original RGB images. The bottom row of each panel shows a boxplot of IoU score for lesions and F1 score for pycnidia.

Evaluating the validation dataset with respect to the annotated and predicted number of pycnidia and rust pustules showed that the annotated and predicted values are highly correlated (see Figure 5). Unfortunately, the range in terms of the numbers of pycnidia and rust pustules differed for direct and diffuse lighting conditions. Nonetheless, in the overlapping regions no significant differences were observed between the performance under diffuse and direct lighting. The same behaviour was observed with PLACL with the exception of multiple outliers under direct lighting where the proposed method overestimated the PLACL values.

**Figure 4:**
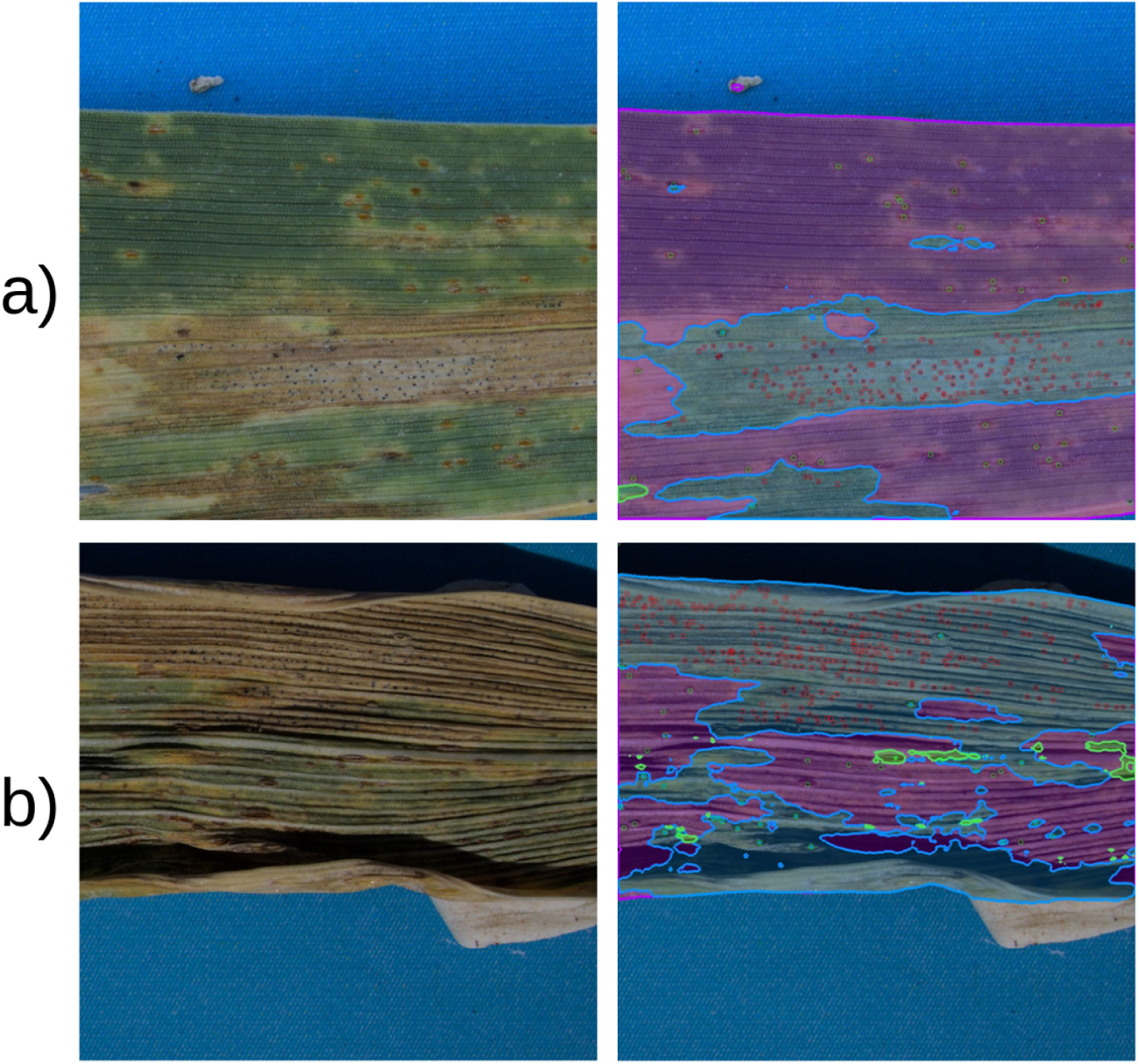
Sample images from EFD dataset and corresponding predictions. Blue masks denote necrotic lesions, purple masks denote leaf and green masks denote insect damage. Red circles denote pycnidia and green circles denote rust pustules. Transparent areas denote the background and regions where the confidence of predictions is below 0.5. Panel a) showcases a success scenario where the method achieves a good performance on all classes. Panel b) demonstrates the performance on an edge scenario with complex lighting resulting in hard shadows and directly illuminated regions at the same time.

**Figure 5:**
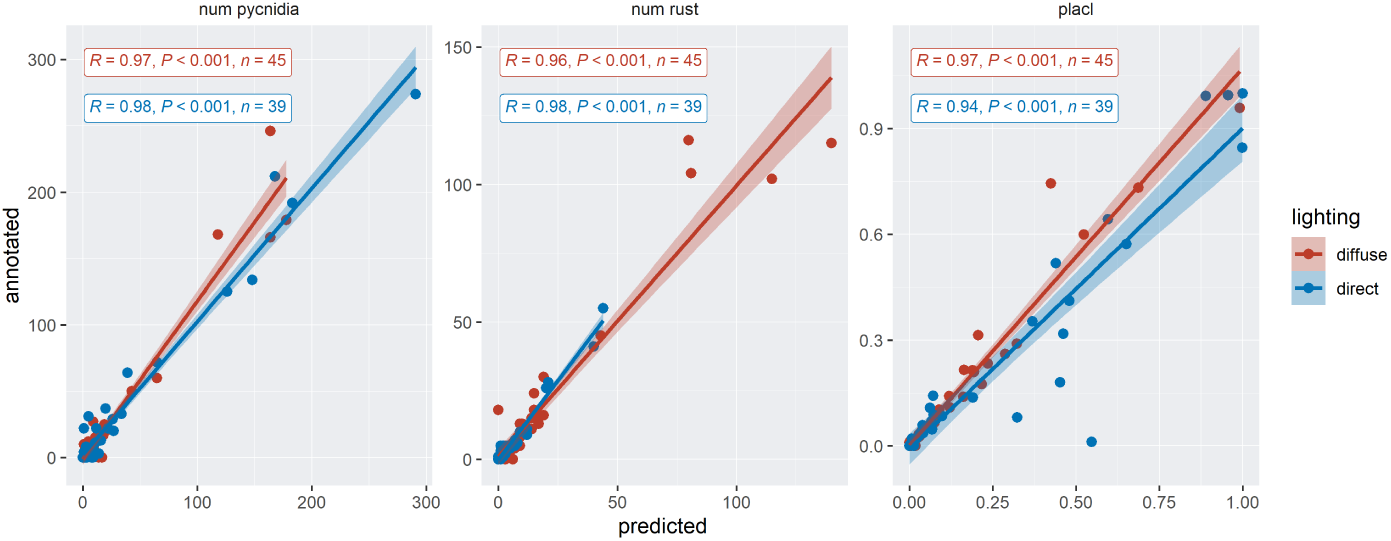
Computed values for number of pycnidia (left panel), number of rust pustules (middle panel) and PLACL (right panel) for each individual image of the validation dataset. Datapoints from images under diffuse lighting are denoted in red whereas images under direct light are denoted in blue.

### 4.3 Generalization Capabilities

#### 4.3.1 Flatbed Scanners Compatibility

Automated image analysis of STB symptoms with flatbed scanners has been the dominant method of symptoms quantification until now. Thus, in the following experiment, images from flatbed scanners were predicted with models trained on the EFD dataset to demonstrate the backwards compatibility of the proposed method. The predictions on images from flatbed scanners are shown in Figure 6. Generally, the major regions are predicted accurately, but the boundaries of individual classes are less smooth and more prone to error (see bottom leaf border of row b) in Figure 6). The already worse performing class of insect damage (see Section 4.1) seems to continue this trend as it is misclassified as leaf in the image row a) in Figure 6. On the other hand, the detection of pycnidia reaches a comparable performance, managing to correctly detect pycnidia even outside of lesions (see row a) in Figure 6). Generally, the proposed method proved to be applicable on images from flatbed scanners, rendering it backwards compatible with existing datasets acquired according to the protocol from Stewart et al. [45].

**Figure 6:**
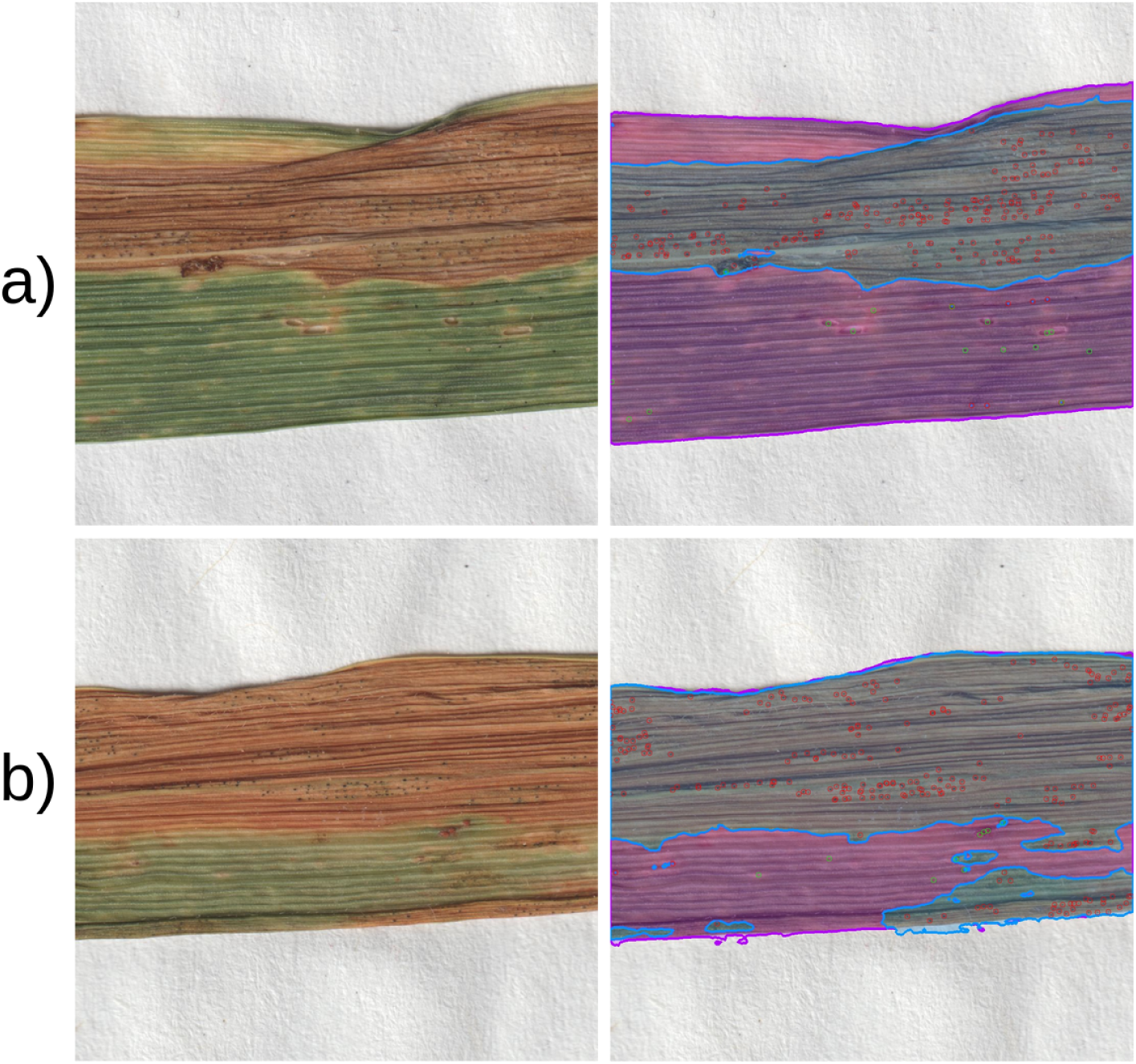
Sample images from flatbed scanners following the protocol of Stewart et al. [45] and the corresponding predictions. Blue masks denote necrotic lesions, purple masks denote leaf and green masks denote insect damage. Red circles denote pycnidia and green circles denote rust pustules. Transparent areas denote the background and regions where the confidence of predictions is below 0.5.

#### 4.3.2 Inference in the Wild

In order to assess the performance in the target environment, the models trained on the proposed dataset were applied to images taken directly in the field without interacting with the plants. We analyzed the performance of the proposed method in regions which corresponded to the appropriate regions of interest with sufficient quality. Within these regions, many additional challenges were present, including different orientations, more complex lighting, backlit leaves, continuous scaling of objects due to the changes in perspective and varying degrees of blur. Under closer examination, necrotic lesions that are in focus are predicted properly (see cyan and red rectangles in Figure 7), however with the decreasing image quality due to factors such as focus blur or direct lighting the prediction quality decreases to favor an incorrect assignment into the background or insect damage classes (see blurred regions of cyan and red rectangles in Figure 7).

A similar behaviour was observed for pycnidia predictions. Areas with high image quality generally show better performance (see cyan rectangle in Figure 7.a), though some areas showed dysfunctional performance in spite of sufficient image quality (see cyan rectangle in row b) in Figure 7). In this case the entire region yielded poor pycnidia prediction. More interestingly, other areas within the same image did yield correct pycnidia predictions associated with them (see red rectangle in row b) in Figure 7). Up- and down-scaling the image as well as lowering the confidence threshold for predictions did not change the poor performance for pycnidia in this region. Besides the aforementioned aspects, the authors could not identify an objective image quality factor that would cause this behaviour and therefore suspect insufficient representation of such a scenario in the EFD dataset.

**Figure 7:**
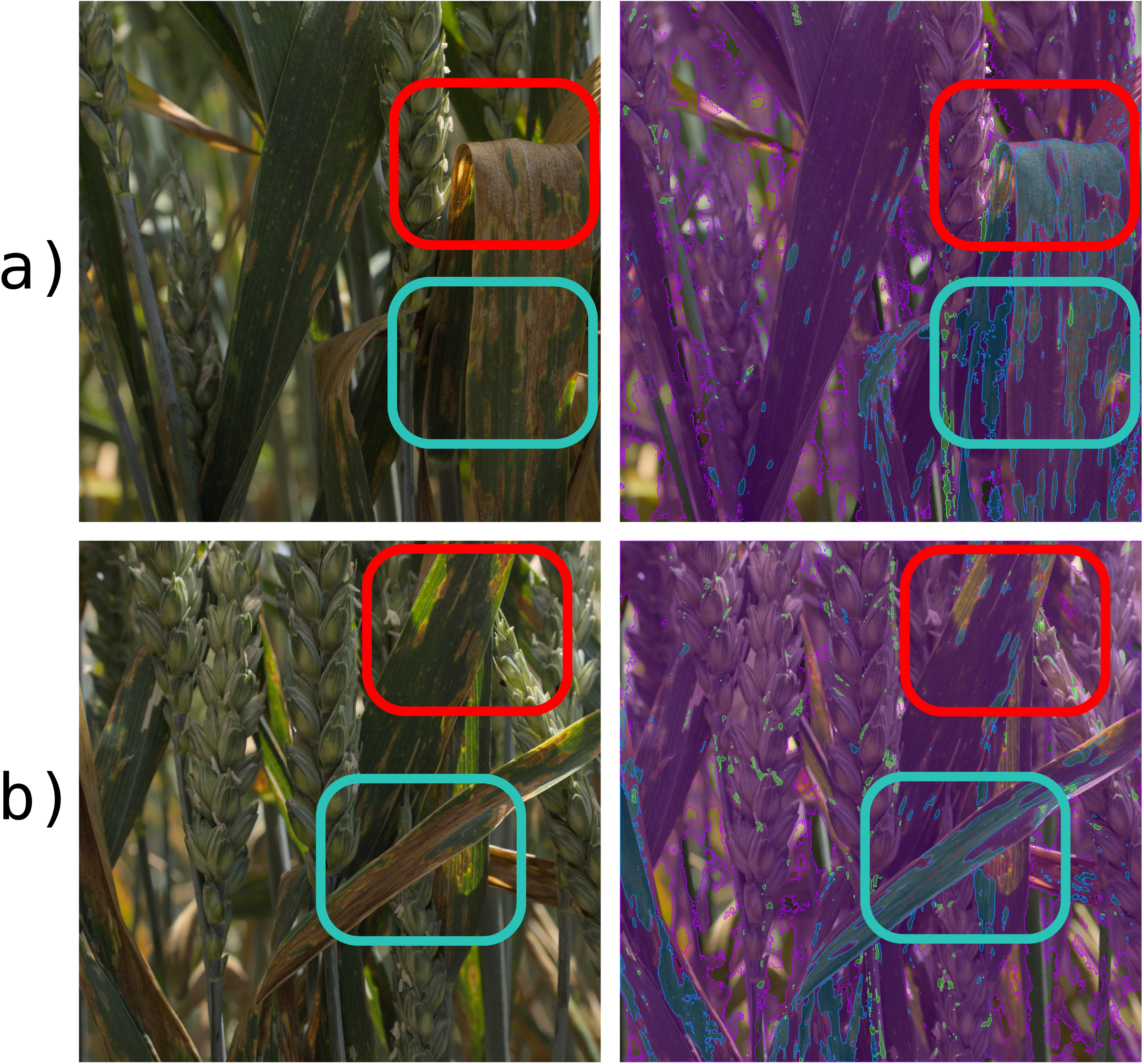
Two images of unstructured canopy directly from the field with their corresponding predictions. The red and cyan rectangles denote specific areas for closer observation. Zoom into the image for sufficient scaling. Blue masks denote necrotic lesions, purple masks denote leaf and green masks denote insect damage. Red circles denote pycnidia and green circles denote rust pustules. Transparent areas denote the background and regions where the confidence of predictions is below 0.5.

## 5 Discussion

We demonstrated a feasible approach to imaging detached leaves at a submillimeter resolution in an uncontrolled outdoor environment. We achieved sufficient resolution to allow accurate imaging of extreme features such as pycnidia which are one of the smallest symptoms on the foliar disease spectrum. The approach of imaging detached leaves outdoors introduces new sources of variance in the form of more variable lighting intensity and potential blur due to a shallow depth of field and potentially curling leaves compared to existing STB evaluation methods which utilize leaves which were pressed flat [45] [34]. The proposed imaging approach allows for imaging of freshly detached leaves, avoiding visual changes in symptoms appearance often caused by the delay between leaf sampling and leaf imaging. However, the proposed imaging approach does not yield any significant benefit in terms of throughput as both the flatbed scanner imaging and the proposed method are bottlenecked by the time needed to identify and detach infected leaves and manipulation of the detached leaves prior to imaging.

The collected images and resulting annotated dataset created an authentic set of disease symptoms under natural field conditions. In contrast to the more common experiments conducted on seedlings that are inoculated with a high concentration of *Z. tritici* spores under highly controlled environmental conditions, the EFD dataset provides a much more representative picture of the real world complexity needed to develop applications that will be useful for breeders and farmers. The additional symptoms variance comes from including different host-pathogen interactions associated with the selected sets of cultivars, a set of different *Z. tritici* strains, as well as naturally occurring co-infections with brown rust and insect damage.

Despite the great complexity of the task and the challenging size of the EFD dataset, we were able to deliver a proof of concept for field STB imaging and quantification by utilizing deep neural networks. Within the scope of model families, the hyperparameter search delivered rather shallow variants of the respective models (see Section 4.1) which can indicate that deeper models are too complex for the underlying dataset, while more shallow models offer a way of regularization.

Our proposed method achieves a similar segmentation performance for lesions *F* 1*_necrosis_* = 0.87 as a similar deep learning based method for STB evaluation *F* 1*_necrosis_* = 0.9 [34] in spite of including much more diverse imaging conditions. However, we achieved significantly higher performance on pycnidia detection *F* 1*_pycnidia_* = 0.76 compared to *F* 1*_pycnidia_* = 0.36 [34]. We speculate that this might be due to the older, anchor-based architecture of YOLOv5 [28] in combination with high resolution images with large aspect ratio and small objects to be detected.

We argue that color based methods for STB analysis such as [45] [30] are incapable of conducting the same task of analysis under uncontrolled lighting because they cannot inherently adjust their decision thresholds when conditions change dynamically.

Furthermore, other methods focusing on the quantification of necrosis [48] [8] [11] [52] [3] [6] can be successful in quantifying the necrotic material but due to their lower resolution cannot keep up with the diagnosis aspect, while the high resolution approach can not only detect but also diagnose the damage.

Mathieu et al. [34] concluded that a small dataset was sufficient to deliver a solid performance on their task under controlled conditions. We believe that for the task of field foliar disease analysis, it is essential to provide a large dataset to reach a robust performance because many more scenarios are present when conducting analyses in the field. This can be observed in the disparity of performance on different segmentation classes (see Table 1) probably due to the insufficient representation of the insect damage class. The same reasoning can be applied based on the samples from section 4.2.1 and section 4.3.2 where an image region shows much worse performance for pycnidia detection. This indicates the need to provide more training samples in order to reach a stable performance. Anderegg et al. [7] were able to accurately track disease development in field-based image time series of leaves after extending the dataset by about 10%. This data set represented another season, different genotypes, multiple leaf layers, and contrasting phenological stages. This further underlined the robustness, flexibility, and adaptability of the proposed method.

The method proposed here is compatible with expanding the dataset to improve its performance as well as potentially introducing additional diseases. A larger dataset would unlock the potential for a deeper network that is capable of extracting more complex features from the data and thus being able to achieve superior performance.

Despite its many limitations, the proposed method at its current stage has already achieved a satisfying performance in the context of quantifying STB symptoms (see section 4.2.1). We believe that the next milestone in the development of a disease quantification tool will be to improve the throughput and minimize the necessary human input. Since the leaf sampling and manipulation requires the greatest time investment, the returns of faster inference are diminishing (see section 4.2.1) and the method would greatly benefit from being able to avoid direct sampling of individual leaves.

### 5.1 In Field Analysis

A key element to develop a high throughput evaluation pipeline is to make images directly from crops in the field under natural conditions. Not only will this eliminate the need for time consuming manual manipulation of individual leaves but it will also render the analyses non-invasive [27]. The preliminary requirement to achieve this goal is the ability to analyze images collected under diverse lighting conditions. The work described here has shown that the state of the art computer vision is indeed capable of fine grained analysis of foliar diseases under natural light. Furthermore, experiments on field images (see section 4.3.2) show that the analysis in this highly unstructured scene is indeed possible, though it will require additional steps which should be the focus of future research and development. The key missing capability is to robustly identify the correct reference region that will be used to collect data. This identification will depend on a mixture of parameters based on the goal of the analysis. For STB the reference region should include parts of an image which have sufficient quality to analyze the symptoms and especially identify pycnidia whilst identifying leaf surfaces that are in the correct location within the plant morphological structure.

With the appropriate dataset, such a pipeline will be capable of quantifying not only STB and brown rust but also other common wheat diseases (e.g. yellow rust, powdery mildew, and fusarium head blight) and thus offer a powerful new tool for plant breeders to conduct high-throughput screening of symptoms associated with the most common cereal diseases.

In addition, such approaches will allow not only for disease detection and quantification but also for assessment of specific symptom phenotypes which can provide additional insights into pathogen-host interactions [46] [24] [32] [5].

### 5.2 Outlook

As indicated in Section 7, the proposed method trained solely on the EFD dataset is already capable of a solid performance in some regions on field images with unstructured canopies. In order to achieve a more robust performance, the training dataset will need to be extended by annotating samples containing new scenarios, particularly including more complex lighting and partial blur. Currently, the proposed method is incapable of selecting the appropriate regions where the disease analysis should be conducted. Fortunately selecting the appropriate regions prior to the symptoms analysis can be divided into separate problems, including removing the out-of-focus regions, and recognizing wheat leaves and the general background. The former can be achieved by abstracting the task to focus estimation, depth estimation or texture analysis which are disciplines of active research in the computer vision community. The latter is a task of plant organs segmentation which was already applied in the context of agriculture and foliar diseases [8] [4] [39].

Once these additional processing steps are available, the proposed method will become arbitrarily scalable, since no direct interaction with plants will be required for data acquisition. In addition, this allows in-field data acquisition to be standardized and potential operator-induced measurement bias to be eliminated by employing autonomous agricultural machines.

## 6 Conclusion

The proposed method describes a novel approach for data-driven sub-millimeter analysis of foliar diseases on detached wheat leaves under outdoor conditions. Using the case study of STB we demonstrate the ability of the method to perform in complex and ambiguous scenarios induced by varying external factors including lighting, visually different wheat cultivars and co-infection with other diseases. The already solid performance of the proposed method can be further improved by extending the datasets to include new samples and allows for further extensions to additional new diseases. The proposed method lays the groundwork for future applications in field conditions with the goal of eliminating the need for manual sampling and manipulation of leaves. Combining the proposed method with focus estimation, organ segmentation and automation of data acquisition will unlock its full potential for resistance breeding and disease monitoring and management.

## 7 Acknowledgements

We express our gratitude to J. Alassimone for providing guidance and handson support in preparing the artificial pathogen inoculations. We also extend our appreciation to the Group of Crop Science at ETH Zürich, particularly S. Corrado for their expertise in crop husbandry and B. Herzog for their assistance in seed preparation and management. We also would like to thank S. Vuillemin, S. Gürkan, K. Gefe, and T. Khampo for assistance with image acquisition and annotation.

## 8 Data availability

The datasets used in this study are available from the corresponding author on reasonable request. Trained models and code for inference are available under: https://github.com/RadekZenkl/leaf-toolkit

## Footnotes

1 https://docs.ultralytics.com/datasets/pose/

