## Supplementary Table A1 for "Towards high throughput in-field detection and quantification of wheat foliar diseases with deep learning"

### A Appendix

| Predicted \ True | Pycnidium | Rust | Background |
|---|---|---|---|
| Pycnidium | 1569 | 0 | 401 |
| Rust | 0 | 711 | 163 |
| Background | 909 | 303 | - |

Table A1: Confusion matrix for keypoint detections
